# Lactate Transport and Signaling Mediated by AMD3100 Ameliorates Astrocyte Pathology and Remyelination Without Additional Extension of SOD1^G93A^ Mice’ Life-Span

**DOI:** 10.1101/2022.01.28.478264

**Authors:** Inna Rabinovich-Nikitin, Beka Solomon

## Abstract

**Background:** Amyotrophic lateral sclerosis (ALS) is characterized by the progressive degeneration of motor neurons accompanied by the accumulation of the morphologically and functionally altered glial cells. Reactive astrocytes and microglia secrete pro-inflammatory mediators and contribute to disease progression. Notably, oligodendrocyte functions are disrupted in ALS, including trophic support, myelination, and oligodendrocyte differentiation. ALS patients and mutant SOD1 mice models display reduced levels of myelin basic protein (MBP) and monocarboxylate transporter 1 (MCT1) contributing to impairment of oligodendrocyte function. Lactate, an active metabolite capable of moving into or out of cells, acts as a signaling molecule, transported exclusively by monocarboxylate transporters (MCTs). We showed previously, that AMD3100 increases the lactate transporter MCT1 in an animal models of Alzheimer’s disease (AD) and ALS.

**Methods and Results:** AMD3100 is a reversible antagonist of CXCR4, and therefore inhibits the CXCR4/CXCL12 axis. AMD3100 was shown to have beneficial effects on extension of SOD1^G93A^ mice’ life-span and enabling migration of hematopoietic stem cells (HSPCs) from bone marrow to periphery. The low content of lactate and transporters in SOD1^G93A^ mice model led us to propose a combined treatment with AMD3100 and exogeneous L-lactate.

**Conclusions:** The combined treatment attenuated neuroinflammation and remyelination but did not have a significant extension of SOD1^G93A^ mice’ life-span compared with AMD3100 treatment alone.

## Introduction

Amyotrophic Lateral Sclerosis (ALS) disease is caused by many interactions between molecular and genetic pathways. Mitochondrial dysfunction, oxidative damage, misfolded proteins, defective axonal transport, excitotoxicity and neuroinflammation, all contribute to the progression of the disease [1]. Pathological findings in motors neurons (MN)s reveal evidence of a significant involvement of non-neural cell types, such as the reactive astrocytes and activated microglia, which secrete neurotoxic factors and pro-inflammatory cytokines [1].

Microglia, the immune cells within the CNS, can either have a neuroprotective or a neurotoxic role during ALS progression. It is believed that microglia protect neurons early in the disease stage however, as the disease progresses, activated microglia can be pro-inflammatory and neurotoxic [2, 3]. Interestingly, in the mutant SOD1 (mSOD1) animal model, removing the mSOD1 from astrocytes postponed the activation of microglia and, therefore, increased survival [2].

Astrocytes affect oligodendrocytes by creating an environment that promotes oligodendrocyte progenitor cells (OPCs) recruitment and their migration and differentiation [4]. Dysfunction of oligodendroglia leads to axon degeneration and is depended on disruption of energy metabolites such as lactate or glucose [5]. Lactate ,an active metabolite ,is transported exclusively by monocarboxylate transporters (MCTs), and changes to these transporters alter lactate production and use [5]. Recent publications suggested a role for oligodendroglia MCT1 in the astrocyte-neuron lactate transport, which is an important source of trophic support to neurons. Lactate is a major carbon source fueling metabolic pathways and a key molecule regulating diverse biological processes. Maintaining homeostatic circulating levels of lactate may ameliorate and even reverse neurodegenerative disease progression. Recent data showed that the low content of cerebral lactate and lactate transporters may lead to blockage of lactate transport from glia to neurons and therefore, alternate lactate metabolism in AD brains [6].

Using lactate as an energy fuel for neurons in neurodegenerative diseases, such as ALS and AD, may improve symptoms by increasing MCT1. Notably, however, as recently reported [7] no significant delay in age of disease onset or in the median survival or disease duration of SOD1^G93A^ mice was reported after the treatment with a viral vector encoding MCT1. The treatment up-regulated MCT1 lactate transporter in white matter oligodendrocytes, but did not ameliorate the disease outcome in the mice. These findings were explained by the observation that local MCT1-mediated lactate delivery failed to reach the extremely long axons in the white matter or that properly functioning MCT1 transporters might be compromised by defects in downstream lactate transport, accumulating in the periaxonal space due to lack of axonal uptake or lack of lactate. Based on this, we propose here lactate supplementation in addition of increase of MCT1 as a treatment to increase neuronal survival. Lactate is required to sustain contraction of skeletal muscle. Administration of exogenous L-lactate may increase the available lactate to nerves and neurons and thus improve the symptoms of ALS. Neurons and peripheral nerves are dependent on glial cells and oligodendrocytes respectively to support their high energy demands by shuttling lactate to neurons and peripheral nerves We previously demonstrated that chronic administration of AMD3100, a reversible antagonist of CXCR4 to SOD1^G93A^ mice led to significant extension of mouse lifespan, improved motor function and increase in the MCT1 transporters [8]. Combined administration of both AMD3100 and exogenous L-lactate to SOD1^G93A^ mice will enable the utilization of endogenous and exogeneous lactate, enhance myelination and attenuate neuroinflammation.

## Materials and methods

### Transgenic mice and treatment

Transgenic mouse strain expressing mutant human SOD1 with the ALS-causing mutations: hemizygous SOD1^G93A^ was used in this study. SOD1^G93A^ mice were maintained on a C57BL/6 congenic background. 50 days old female SOD1^G93A^ mice were treated subcutaneously with 5mg AMD3100 (Sigma-Aldrich, USA) or 5mg AMD3100+1mM lactate or 1mM lactate (Sigma-Aldrich, USA) or PBS (Biological Industries, Israel) twice a week. AMD3100 and lactate do not create a chemical binding in a tube. During treatment, body weight, motor functions and survival rate were examined. For biochemical analysis, mice were sacrificed at 110 days old via i.p anesthesia administration of 100 mg/kg Ketamine (Fort Dodge, USA) and 20mg/kg Xylazine (Merck, Germany) following trans-cardial perfusion with saline. The spinal cords were collected and processed as described below.

The animals were in house maintained colony. Mice were housed in standard conditions: constant temperature (22±1°C), humidity (relative, 40%), and a 12-h light/dark cycle and were allowed free access to food and water. All of the animal experiments were conducted in accordance with the Guide for the Care and Use of Laboratory Animals and were approved by the Institutional Animal Care and Use Committee of Tel Aviv University.

### Rotarod test

The Rotarod test was performed once a week at the same approximate hour and before treatment to decrease fluctuations in performances of mice that might occur due to stress or difference in awareness according to the mice daily cycle. Each mouse was placed on a textured drum (NBT biotech) rotating at an increasing speed ranging from 4-40 RPM, with a constant acceleration of 1RPM per 10sec. The mice were trained on the machine for three days before the actual beginning of the analysis. Mice were then tested once a week, three attempts, for the length of the experiment. The time each mouse remained on the drum was recorded, up to 300sec.

### Survival analysis

Mice were examined each day, and date of death was recorded. Mice received treatment until the end stage of the disease, which was defined as the point at which animals could not right themselves within 30 sec after being placed on their side. At that point mice were euthanatized with Co2. In addition, mice which exhibited extreme signs of distress or which lost over forty percent of initial body weight were sacrificed.

### Determination of disease onset

Disease onset was determined as the time animals reached their maximum bodyweight.

### Tissue fractionation

SOD1^G93A^ mice treated with AMD3100 at 50 days old were sacrificed at the age of 110 days and their spinal cords were collected. Tissues were homogenized on ice in 5 volumes (w/v) of T-per extraction buffer (Pierce, USA) complemented with protease inhibitor tablets (Complete Mini Protease Inhibitor Tablets, Roche) and phosphatase inhibitor cocktail tablets (phosSTOP, Roche). After sonication, the homogenates were centrifuged at 10,0000g for 1h at 4°C. For muscle fraction, the gastrocnemius muscle was collected, homogenized on ice in 5 volumes (w/v) of T-per extraction buffer complemented with protease inhibitor tablets (Complete Mini Protease Inhibitor Tablets, Roche) and phosphatase inhibitor cocktail tablets (phosSTOP, Roche), 0.5% triton-100, 1% sodium deoxycholate and 3% SDS. After sonication, the homogenates were centrifuged at 10,0000g for 1h at 4°C. The resulting supernatants represented the whole muscle fraction. Protein concentrations were determined using BCA protein assay kit (Thermo, USA).

### Western immunoblot analysis

Equal amounts of mice spinal cord homogenate protein (40 μg) were resolved on SDS-PAGE, transferred to nitrocellulose membrane and blocked overnight with 5% skim milk in TBS-T (0.3% Tween 20). Blots were probed with the following primary antibodies: mouse anti Actin (1:10,000 Sigma-Aldrich, USA), mouse anti Myelin basic protein (MBP) (1:1000 Covance, USA), mouse anti BACE1 (1:1000 Millipore, Germany), mouse anti MCT1 (1:1000, Santa cruz, USA), rabbit anti GFAP (1:1000, Dako, USA), mouse anti S100B (1:500, Novus Biologicals, USA), rabbit anti-IL-6 (1:1000 Peprotech, USA), and mouse anti-Iba-1 (1:1000 Millipore, Germany). Blots were incubated with anti-mouse or anti-rabbit secondary antibodies conjugated to peroxidase (Sigma-Aldrich, USA) and developed with the EZ-ECL detection kit (Biological Industries, Israel). Quantitative densitometric analysis was performed using the densitometric software EZQuant-Gel (version 2.12).

### Histological staining

A separate cohort of SOD1G93A mice treated with AMD3100 (n=3) or PBS (n=3) at 50 days old were sacrificed at 110 days old by i.p anesthesia with ketamine/xylazine and perfusion with saline. Their spinal cords were harvested, fixed in 4% (w/v) paraformaldehyde (PFA) in PBS (pH 7.4) and cryoprotected in 30% sucrose in PBS. 25 μm free floating cryosections were prepared from lumber spinal cords and stained for Luxol fast blue stain for myelin. 15 sections per group were used. Sections were de-fatted for 30 min in 1:1 chloroform/ethanol, rinsed in 95% ethanol and stained with 0.1% (w/v) luxol blue for 12h at 56oC. Excess staining was rinsed in ethanol and differentiated in 0.05% lithium carbonate following 70% ethanol. Sections were dehydrated in graded alcohol, cleared in xylene and coverslipped with enthelan (Merck, Germany). Images were captured by a CCD color video camera (ProgRes C14, Jenoptic, Jena, Germany) attached to a Leica DMLB microscope (Leica, Germany) and analyzed with Image-J Software (NIH, freeware).

### Statistical analysis

All data presented as the mean ± SEM, and subjected to unpaired one-tailed Student’s t-tests. *p<0.05 was considered statistically significant.

## Results

### AMD3100 increases the levels of monocarboxylate transporter 1 of lactate

Lactate is transported exclusively by monocarboxylate transporters (MCTs), and reduction of these transporters affects lactate production and consumption [5, 9]. In the CNS, the most abundant lactate transporter is monocarboxylate transporter 1 (MCT1), which is highly enriched within oligodendroglia, but is also found in muscles [5, 10]. Disruption of MCT1was evident in early stages of ALS in both patients and mouse models, leading to axon damage and neuronal loss [9]. In figure 1 we show that AMD3100 treatment increases MCT1 levels not only in spinal cords (Figure 1A), but also in muscles (Figure 1B) of 50 days old SOD1^G93A^ mice. Interestingly, LM age matched mice treated with AMD3100 also showed significant increase in MCT1 levels in spinal cords (Figure 1C), suggesting AMD3100’s role in preservation of MCT1 expression, with a potential clinical relevance in other diseases as well.

**Figure 1.**
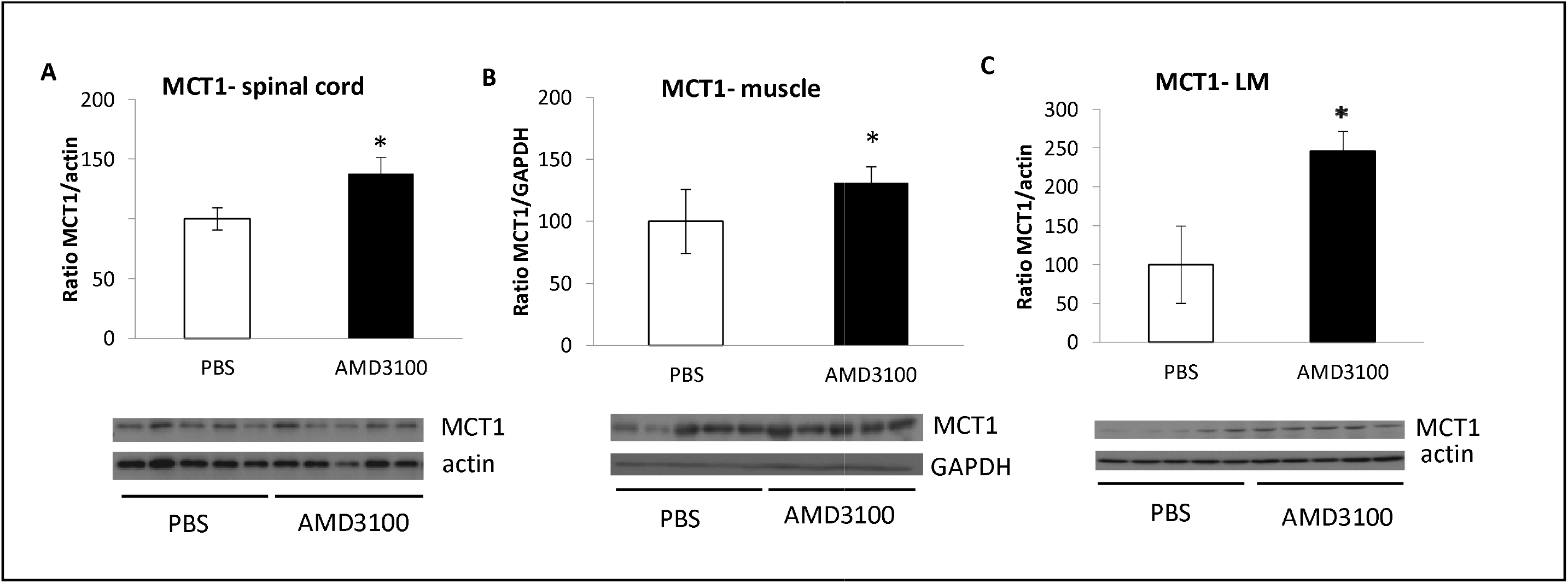
AMD3100 treatment increases MCT1 expression. 50 days old SOD1^G93A^ or LM mice were treated with AMD3100 (n=5) or PBS (n=5), sacrificed at 110 days old and their spinal cords and muscles were analyzed for MCT1 levels. A. MCT1 levels in spinal cord. B. MCT1 levels in muscles. C. MCT1 levels in LM mice. Results are mean ± S.E.M, T-test; *p<0.05.

### AMD3100+lactate treatment does not affect survival and well-being of SOD1^G93A^mice

In order to examine the effects of AMD3100+lactate in vivo, female SOD1^G93A^ mice were treated subcutaneously with PBS (n=4), AMD3100 (n=3), AMD3100+ lactate (n=5) and lactate (n=4) twice a week. Mice received continuous treatment until the end stage of disease which was defined as the point at which animals could not right themselves within 30 sec, after being placed on their side. Treatment of female SOD1^G93A^ mice with AMD3100+lactate or with lactate alone had no beneficial effect on mice survival (Figure 2A), disease onset (Figure 2B) and weight change (Figure 2C) compared with AMD3100 treatment alone. Both treatments of AMD3100+lactate and lactate alone had similar improvement in survival, disease onset and weight change compared to AMD3100 alone. Importantly, all three treatments (AMD3100, AMD3100+ lactate, and lactate) had better performance in all these parameters, compared to PBS treated mice. Motor function assessed by Rotarod test (Figure 2D), was noticeably better in AMD3100 treatment alone, compared with PBS. AMD3100+lactate had no improvement in motor function compared with PBS, and lactate treatment alone had worst motor performance, compared with PBS.

**Figure 2:**
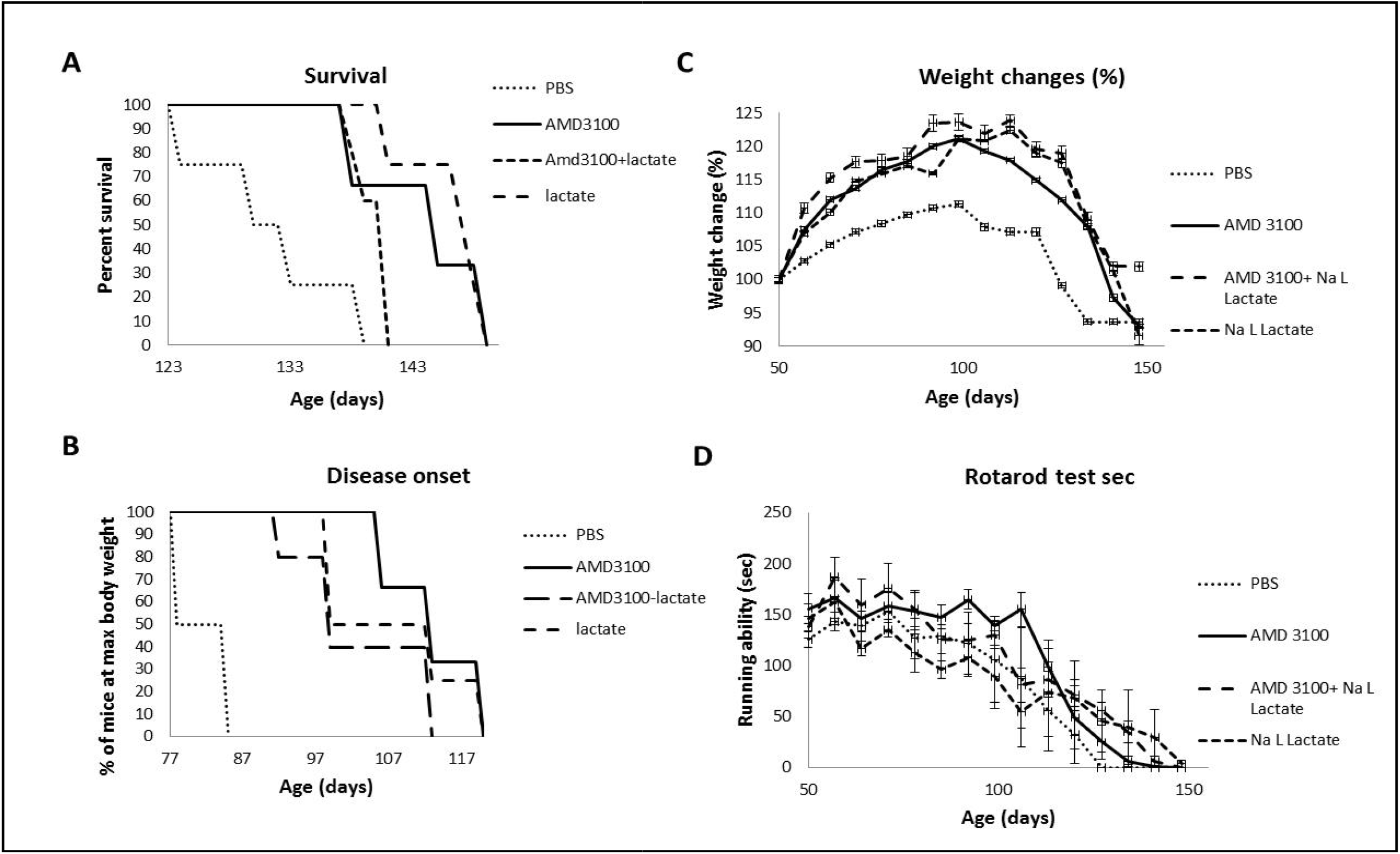
Survival, disease onset, weight changes and motor function plots of SOD1^G93A^ mice treated with AMD3100+lactate. 50 days old SOD1^G93A^ mice were treated subcutaneously with PBS (n=4), AMD3100 (n=3), AMD3100+lactate (n=5) or lactate (n=4) twice a week. A. Survival plots of treated mice. Median survival of PBS= 132, AMD3100= 144, AMD3100+lactate= 141. Lactate= 148. Survival of mice was defined as the point at which animals could not right themselves within 30 sec after being placed on their side. B. Disease onset of treated SOD1^G93A^ mice. Disease onset was determined as the time animals reached their maximum bodyweight. C. Weight changes monitored weekly. D. Motor function assessed weekly by Rotarod test. Results are mean ± S.E.M.

### AMD3100+lactate treatment decreases astrocytic inflammation markers of SOD1^G93A^mice

Lactate measurements in the spinal cord of transgenic mice confirmed that SOD1^G93A^ mice develop metabolic impairment between the age of 30 days, when lactate levels are still normal, and 40 days, when there is a 20% decrease in lactate, with a further decrease by 30% at 60 days, providing an interesting link between the onset of denervation and motor neuron metabolic dysfunction [11]. A recent study showed that lactate supplementation can completely reverse the observed toxicity of astrocytes [12]. Indeed, our data indicate that external supplementation of lactate together with AMD3100 significantly decreased levels of activated astrocytes in 50 days old SOD1^G93A^ mice, as shown by GFAP levels (Figure 3A) and S100B levels (Figure 3B). In contrast, treatment with AMD3100 had no effect on astrocytes activation in the same mouse model, emphasizing the role of lactate on astrocyte activation.

**Figure 3:**
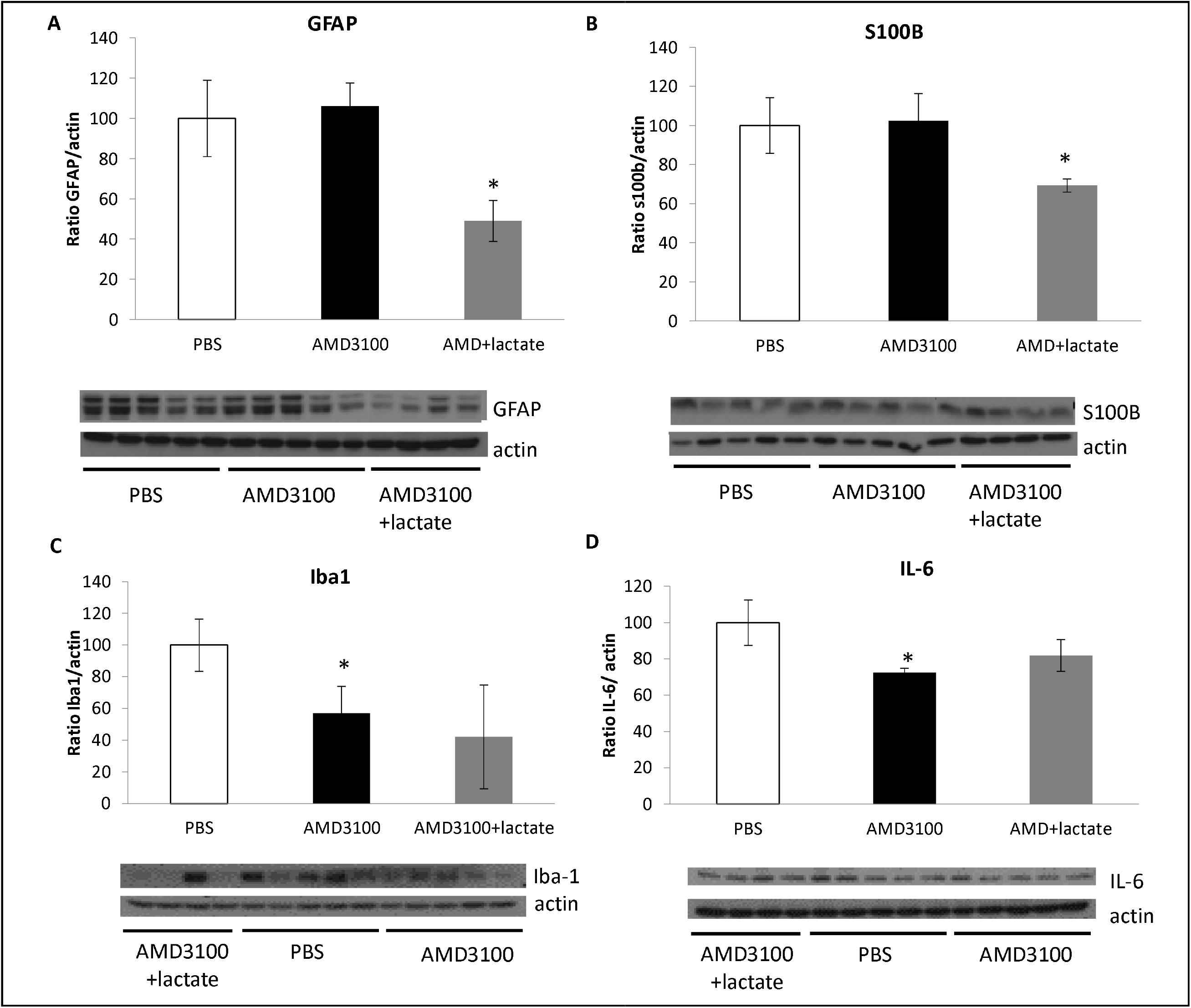
AMD3100+lactate treatment decreases activation of astrocytes but has no effect on microglial markers in SOD1^G93A^ mice. 50 days old SOD1^G93A^ mice were treated subcutaneously with PBS (n=5), AMD3100 (n=5), AMD3100+lactate (n=4) twice a week. Mice were sacrificed at 110 days old and their spinal cords were tested for astrocytic and microglial inflammation markers. A. GFAP levels. B. S100B levels. C. Iba-1 levels. D. IL-6 levels. Results are mean ± S.E.M. T-test; *p<0.05.

### AMD3100+lactate treatment does not affect microglial inflammation of SOD1^G93A^mice

In our model, we found that combined treatment of AMD3100+lactate had no significant effect on the microglial marker Iba-1 (Figure 3C) and on the cytokine IL-6 (Figure 3D). The treatment of AMD3100 alone to 50 days old SOD1^G93A^ mice resulted in significant decrease in those two markers, compared with the AMD3100+lactate treatment that also resulted in some decrease in these markers, however not significantly.

### AMD3100+lactate treatment increases myelin basic protein (MBP) levels, but not BACE1 inSOD1^G93A^mice

AMD3100 was previously shown to increase oligodendrocyte progenitor cells (OPCs) which are responsible for creating the myelin sheets [13]. The current data supports these findings and show that spinal cords of SOD1^G93A^ mice treated with AMD3100 at 50 days old were significantly remyelinated, as shown by western blot analysis for myelin basic protein (MBP) levels (fig. 4A) and by the myelin associated stain, Luxol Fast blue staining, of spinal cord sections (fig. 4B). In our data, the Luxol Fast blue staining revealed no changes in total myelin load between control and AMD3100 treated mice, however, myelin density was significantly increased in the AMD3100 treated group. Furthermore, BACE1 levels, which is known to have a role in myelination [14] were also increased when treated with AMD3100 (Figure 4C).

**Figure 4:**
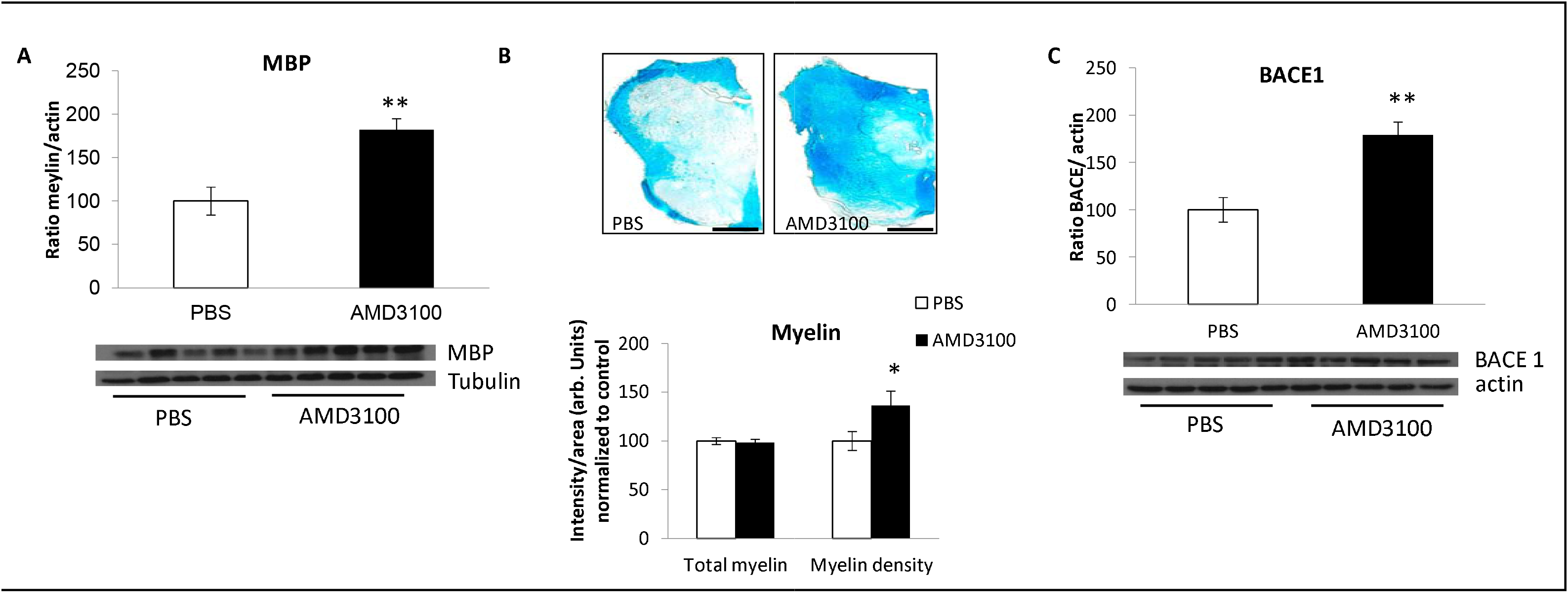
AMD3100 treatment significantly increases MBP and BACE1 in SOD1^G93A^ mice. 50 days old SOD1^G93A^ mice were treated with AMD3100 (n=5) or PBS (n=5), sacrificed at 110 days old and their spinal cords were analyzed for myelin and BACE1 levels. A. Myelin basic protein (MBP) levels. B. 50 days old SOD1^G93A^ mice were treated with AMD3100 (n=3) or PBS (n=3) from 50 days old to 110 days old and stained for Luxol fast blue. Representative Images of stained sections are presented. C. BACE1 levels. The control group is set to 100%. Results are mean ± S.E.M, T-test; *p<0.05, **p<0.01.

Previous studies suggest that overall oligodendrocytes consume more lactate than neurons. The consumed lactate is used to fuel the mitochondria, for lipid synthesis, and to make myelin [15]. Another study showed that oligodendrocytes consume lactate by MCT1. In this study, myelination was rescued when exogenous L-lactate was available for the oligodendrocytes. Furthermore, lactate enhances development and myelination of oligodendrocyte especially when the levels of glucose are low. [16]. In diseases that involve energy deprivation during myelination, lactate transporters in oligodendrocytes may play a pivotal role in rescuing the inhibition of myelination that occurs. In figure 5A we show that 50 days old SOD1^G93A^ mice treated with AMD3100+lactate, significantly increased their myelin basic protein (MBP) levels compared with AMD3100 alone. However, BACE1 levels were not affected following AMD3100+lactate treatment, compared with PBS (Figure 5B).

**Figure 5:**
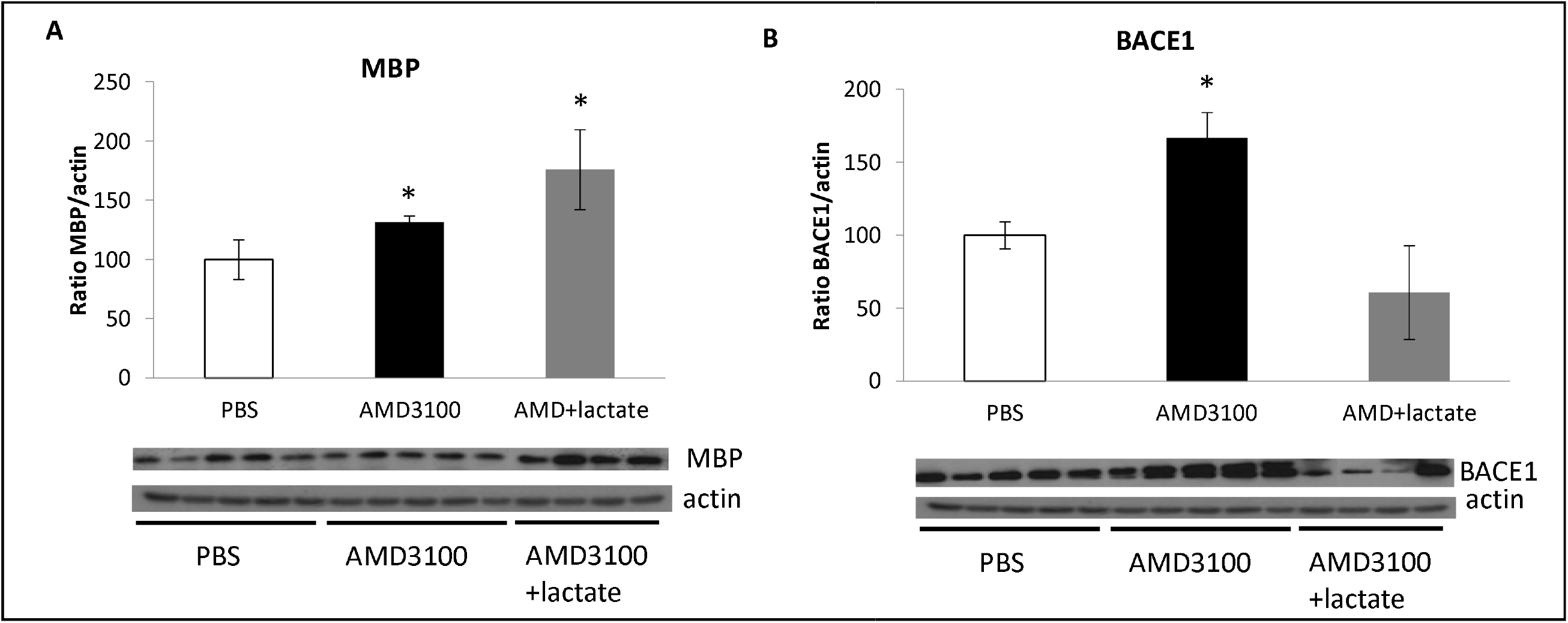
AMD3100+lactate treatment significantly increases MBP but not BACE1 in SOD1^G93A^ mice. 50 days old SOD1^G93A^ mice were treated subcutaneously with PBS (n=5), AMD3100 (n=5), AMD3100+lactate (n=4) twice a week. Mice were sacrificed at 110 days old and their spinal cords were tested for A. MBP levels. B. BACE1 levels. Results are mean ± S.E.M. T-test; *p<0.05.

## Disscusion

Herein, we showed that inhibition of CXCR4/CXCL12 axis by combined treatment of AMD3100+ lactate resulted in increase in MCT1 levels, enhanced myelination and attenuation of astrocytes reactivation without affecting microglia activation.

Notably, treatments of AMD3100+lactate and lactate had similar improvement in survival, disease onset and weight change compared to AMD3100 alone. All three treatments (AMD3100, AMD3100+ lactate, and lactate) exhibit better performance in all these parameters, compared to PBS treated mice (Figure 2). By using the combined approach of AMD3100+ lactate in the SOD1G93A mouse model of ALS, we did not observe a functional rescue of disease phenotype, nor a survival benefit on top of the beneficial effect of AMD3100 alone.

Therefore, we suggest that lack of effect of exogenous lactate can be explained by insufficient amount of MCT1 for transport of exogenous and endogenous lactate. As a consequence, we confirmed the previous reported data that conclude that upregulation of MCT1 lactate transporter in white matter oligodendrocytes does not ameliorate the disease outcome in this ALS model mice [7].

The analysis of spinal cords of SOD1^G93A^ mice treated with AMD3100 and lactate at 50 days old showed increased remyelination, as shown by western blot analysis of MBP levels (fig. 5A) and by the myelin associated stain, Luxol Fast blue, of spinal cord sections (fig. 5B) which revealed the myelin density was significantly increased in the AMD3100 treated group. Moreover, the endopeptidase BACE1, a marker which has a role in regulating the process of myelination and myelin sheath thickness in the central and peripheral nerves, showed significant increase in 50 days old SOD1^G93A^ mice treated with AMD3100 (Figure 5C).

These findings were further supported by the observation that supplementation of external lactate together with AMD3100 significantly decreased levels of activated astrocytes as shown by GFAP levels (Figure 3A) and S100B levels (Figure 3B). In contrast, treatment with AMD3100 alone had no effect on astrocytes activation in the same mouse model, emphasizing the role of lactate on astrocyte activation.

Combined treatment of AMD3100 and lactate did not show additional effect on activated microglia compared to AMD3100 alone. During initial stages of disease, microglia exhibit an M2 phenotype supporting neuronal survival [2]. During initial stages of disease, microglia exhibit an M2 phenotype supporting neuronal survival [2]. Yet, as the disease progresses, microglial activation changes towards an M1 phenotype, i.e. activated neuroinflammatory microglia. This phenotype can then induce neurotoxic reactive astrocytes [17]. Indeed, microglial activation was shown not only during disease onset and progression in mutant SOD1 mice model of ALS but also during later disease stage, suggesting that microglia-targeted therapy may be an effective strategy for ALS treatment.

The CXCR4/CXCL12 (SDF1) axis is one of the major signal transduction cascades involved in the inflammation process and regulation of homing of hematopoietic stem cells (HSCs) within the bone marrow niche [18]. The inhibition approach of CXCR4/CXCL12 signaling by AMD3100, a reversible antagonist of CXCR4, has been shown beneficial in increasing the survival of treated ALS mice in addition to enhancing mobilization of endogeneous hematopoietic stem cells (HSCs) from the bone marrow to periphery [8]. The bone marrow contains hematopoietic stem and progenitor cells HSPCs-potential targets for cell-based therapy. The derived CD34+ HSCs possess a major advantage as compared to other stem cell types, as they are easily mobilized endogenously leading to their blood-borne circulation, to home to injured or inflamed tissues in any compartment of the human body including the CNS [19]. Bone marrow derived –cells (BMDC)s may deliver neuroprotection and neuro-regeneration via various mechanisms, including trans-differentiation or cell-cell fusion with resident cells [19]. Bone marrow cells also benefit the injured tissue by secreting bioactive factors, which can repair and improve neurogenesis, convey intrinsic repair and enhance neurogenesis [18]. Our data propose that beneficial effect of AMD3100 on survival and well-being of treated mice is due to bone marrow-derived microglia-like cells that can enter the CNS under normal physiological conditions and preferentially present in regions suffering from neurodegeneration or exogenous damage. These findings suggest that the disease rises from a damage within MNs and their glial associates, rather than solely from neuronal injury.

Importantly, all three treatments (AMD3100, AMD3100+ lactate, and lactate) had better performance in all these parameters, compared to PBS treated mice. Using the combined approach of AMD3100+ lactate in the SOD1^G93A^ mouse model of ALS, did not contribute to additional functional rescue of disease phenotype, on top of the beneficial effect of AMD3100 alone. As a consequence, we confirmed the previous reported data [7] that concluded that upregulation of MCT1 lactate transporter in white matter oligodendrocytes does not ameliorate the disease outcome in these ALS model mice in spite of the increased remyelination and reduced astrocytes activation.

Contrary to these findings, the combination of AMD3100 and L-lactate was found to have a beneficial effect in acute model of AD by improving cognitive deficit, reducing tau and APP pathologies and most importantly, leading to shift in microglia to anti-inflammatory M2 profile [20]. The observed downregulation of IL-6 ,TNFα, and MCP-1 together with upregulation of IL-4 and IL-10 may explain the anti-inflammatory effect that we attained by shifting the lineage of the microglia. The occurrence of hematopoietic cells in the brain was validated by the abundance of c-KIT and Sca-1. Interestingly, only the combined treatment of AMD3100 and L-lactate increased BDNF and synaptophysin levels, both of which affect cognition directly [20].

## Conclusion

Overall, the multi-layered role of inhibition of the CXCL12/CXCR4 signaling pathway may lead to prevention of neuroinflammation via mobilization of HSCs from the bone marrow to the bloodstream. AMD3100 is an antagonist of the interaction between CXCL12 and its receptor CXCR4 and can mobilize HSCs within hours rather than days. In addition, AMD3100 is reported to be well tolerated, with no significant side effects, as supported by various clinical trials conducted to date [21].

We suggest that AMD3100 can be considered as an alternative approach for the multistep actions of transplantation of stem cells in the treatment of ALS. Our experimental data supports this notion and suggests AMD3100 as a safe and effective mobilizer of endogenous hematopoietic stem cells.

## Acknowledgments

We are thankful to Becki Barbiro for all her technical assiatance with the studies.

## Notes

### Competing Interest Statement

The authors have declared no competing interest.

